# nanoFOLD : sequence design of nanobodies via inverse folding

**DOI:** 10.1101/2025.04.29.651236

**Authors:** Dawid Chomicz, Bartosz Janusz, Sonia Wrobel, Pawel Dudzic, Adithya Polasa, Kyle Martin, Steven Darnell, Stephen R. Comeau, Konrad Krawczyk

## Abstract

Antibodies devoid of light chains are a promising class of biotherapeutics. Computational methods that address these molecules are crucially needed to accelerate the traditional, long and expensive experimental process of their discovery. Inverse folding, wherein one is tasked to predict a sequence given molecular coordinates, is an established method in scaffold-based protein design. Here we develop an inverse folding method specific to nanobodies. We demonstrate its application in nanobody-engineering scenarios of enriching binders from next-generation sequencing experiments and novel binder design.

## 1 Introduction

Antibodies have revolutionized therapeutic approaches due to their ability to specifically target and neutralize pathogens or diseased cells. Their high specificity and binding affinity make them indispensable in treating various conditions, including cancers, autoimmune diseases, and infectious diseases. However, antibodies are large, complex molecules, often requiring sophisticated production and delivery methods. In contrast, nanobodies, a unique subset of antibodies derived from camelids, are single-domain antibody fragments. These smaller, more stable molecules retain the high specificity of conventional antibodies but offer several advantages. Nanobodies can penetrate tissues more effectively, are believed to be less immunogenic than murine antibodies, and can be produced more economically in microbial systems. These features make nanobodies particularly attractive for therapeutic applications where traditional antibodies face limitations, such as in targeting inaccessible antigens or in environments where stability is a concern (Jin et al. 2023).

There are currently 18 nanobodies in clinical use, including four approvals, Caplacizumab, Envafolimab, Ozoralizumab and LCAR-B38M (Hadsund et al. 2024; Jin et al. 2023). Continued therapeutic applications of this format could be accelerated by computational methods. In the field of antibodies, which has more than 100 molecules approved and close to 1000 in the clinic, there are a plethora of computational methods that currently accelerate the development of these molecules (Norman et al. 2020; Wilman et al. 2022). Though cognate to antibodies, nanobodies require format-specific model development as demonstrated with language models (Hadsund et al. 2024; Li et al. 2023), immunogenicity prediction (Ramon et al. 2023; Sang et al. 2021) or structure prediction (Cohen, Halfon, and Schneidman-Duhovny 2022).

A computational method of specific interest to nanobody development is inverse folding. It involves determining the amino acid sequence that will fold into a given protein structure, essentially reversing the traditional approach of predicting structure from sequence (Hsu et al. 17--23 Jul 2022; Dauparas et al. 2022). This problem is pivotal in protein design, as it enables the rational creation of proteins with desired functions and stability by specifying their three-dimensional structures first.

In the context of antibody and nanobody design, solving the inverse folding problem allows researchers to design novel sequences that adopt structures with high affinity and specificity for target antigens (Bennett et al. 2024). Though extensively applied in proteins (Dauparas et al. 2022), the applications of inverse folding to antibodies have been much less spectacular. Antibodies are notoriously difficult to handle computationally, as evidenced by their classification as one of the most challenging protein groups in AlphaFold3 (Abramson et al. 2024).

It was demonstrated that fine-tuning either proteinMPNN (Dreyer et al. 2023) or ESM-IF (Hummer et al. 2023) on antibodies improves performance. To the best of our knowledge there were no such studies addressing nanobodies. To address this gap, here we develop a nanobody-specific inverse folding framework based on ESM-IF and benchmark it on nanobody-specific tasks.

## 2 Methods

### 2.1 Fine tuning procedure

We employed ESM-IF architecture and weights as a basis for fine tuning. Briefly, ESM-IF1 architecture integrates graph-based and transformer-based methodologies to enable protein inverse folding. It consists of four Graph Neural Network Geometric Vector Perceptron (GVP-GNN) layers for processing geometric information and eight generic Transformer encoder and decoder layers for sequence prediction. The model is invariant to rotation and translation of input spatial coordinates, ensuring robustness in protein structure representation. To handle multi-chain antibody structures, the backbone coordinates of the light chain are concatenated to those of the heavy chain with a 10-position padding of “gap” tokens, which are represented as missing coordinates. This enables seamless processing of antibody variable domain structures while leveraging the pre-trained single-chain capabilities of ESM-IF1. The model was trained on a combination of AlphaFold2 models and predicted structures. Side-chain stripped coordinates are given to the model, which is tasked in reconstructing the sequence (Table 1).

**Table 1.** Models employed in this study. We used ESM-IF as the base, protein-generalistic model. We compared against an antibody-specific AntiFold that is also based on ESM-IF. There are three types of nanoFOLD model developed, X-ray only (xal), model-only (model) and model-only fine tuned on X-ray (ft).

| Model Name | Structure Model | Training Models | Comment |
| --- | --- | --- | --- |
| nanoFOLD-xal | Real X-ray structures | 435 crystal structures | ESM-IF fine tuned only on X-ray structures of nanobodies |
| nanoFOLD-model | NanoBuilder2 | 21,276 models | ESM-IF fine-tuned on models of nanobodies |
| nanoFOLD-ft |  | 21,276 models + 435 crystal structures | nanoFOLD fine-tuned on X-ray structure of nanobodies |
| AntiFold | ABodyBuilder2 | ~150k models + ~2000 crystal structures | ESM-IF fine tuned on antibody structures |
| ESM-IF | AlphaFold2 | 12 million models and 16,000 crystal structures | Foundational model trained on protein structures from AlphaFoldDB |

An antibody-specific model, AntiFold (Hummer et al. 2023) was used here for comparison. Similar to ESM-IF, a combination of models using ABodyBuilder2 and experimental structures of paired heavy/light chains was used to train the model (Table 1). AntiFold builds on the ESM-IF1 inverse folding model through a two-phase fine-tuning strategy specifically designed for antibody structures. The model is first fine-tuned on a dataset of 147,458 predicted antibody structures generated from the Observed Antibody Space (OAS (Olsen, Boyles, and Deane 2022; Kovaltsuk et al. 2018)) using ABodyBuilder2 (Abanades et al. 2023). In the second phase, it is fine-tuned on 2,074 experimentally solved antibody structures from SAbDab (Dunbar et al. 2014), using early stopping to avoid overfitting. To improve sequence recovery, several masking strategies are applied, including shotgun masking (randomly selected positions), span masking (masking consecutive stretches of positions), and IMGT-weighted masking (biasing towards complementarity-determining regions). Gaussian noise is added to backbone coordinates during training to enhance robustness, and layer-wise learning rate decay is applied to preserve weights in earlier layers.

Our fine-tuning of ESM-IF on nanobodies, termed nanoFOLD, comes in three types, following the original ESM-IF and AntiFold efforts to train on . We created 21,276 NanoBodyBuilder2 (Abanades et al. 2023) models, out of a total of 10 million in INDI (Deszyński et al. 2022). We also identified 435 non-redundant (90% sequence identity) structures of nanobodies in the PDB. We trained a crystal-structure only model nanoFOLD-xal and model-only nanoFOLD-model (Table 1). The fine-tuned variety involved further fine-tuning the nanoFOLD-model on the crystal structure dataset. The use of the three models was to reveal any biases/advantages of using the real distribution set (crystal structures), larger but somewhat out-of-distribution datasets (models) and combining the both (fine-tuning).

The fine tuning procedure involved an ADAM optimized, cross entropy with logits loss and a learning rate of 10^-4^ with weight decay of 10^-3^. The parameters were obtained by studying the training procedure of similar models. Firstly, the purposefully small model training dataset allowed for small-scale experimentation with masking regimes and learning schedules, however we did not see a radical benefit of different regimens. Secondly, we were conscious of catastrophic forgetting and we assumed that fine tuning the models on several million nanobody models would inevitably pull the weights away from the local minima obtained for general proteins. Though we note that for specific cases of only CDR-H3 redesign it might be beneficial to re-train an H3-only model, keeping the rest of the sequence intact. Nonetheless, the purpose of this study was to establish whether feeding the model nanobody-specific data, would result in a better performance on a range of realistic biologics discovery tasks.

### 2.2 NGS datasets for enrichment

In our NGS-enrichment experiments we test whether inverse folding can identify sequences with higher propensity for binding against a specific target as it was demonstrated in a similar experiments with antibodies using ESM (Hie et al. 2023). We found two pieces of work that had suitable data, the AVIDA-IL6 and AVIDA-SARS-COV-2 studies (Tsuruta et al. 2023, 2024).

In both studies, nanobodies were raised against a specific immunogen, (IL6 and SARS-COV-2 respectively) and plentiful numbers of binders and non-binders were identified. In each case the immunogens were subjected to mutations, simulating having many more than just a single antigen, though still within a single mutation away.

We only kept the unique sequences for each of the mutants (Tables 2, 3). In the IL-6 dataset we found 650 sequences that were assigned as binders/nonbinders across different mutants. In Sars-cov-2 we found 427 sequences that were assigned as binders/non binders across different mutants. In neither case were there inconsistencies within any antigen which would be problematic for model training.

**Table 2.** Sars-cov-2 dataset statistics. We only kept the sequences that were unique.

| **Mutant** | **Negatives** | **Positives** |
| --- | --- | --- |
| Alpha | 4048 | 1777 |
| Alpha+E484K | 3065 | 1730 |
| Alpha+K417N | 3263 | 1898 |
| Beta | 6473 | 2820 |
| D614G | 6060 | 2353 |
| Delta | 4774 | 1082 |
| Kappa | 1364 | 1284 |
| Lambda | 2591 | 1448 |
| OC43 | 2984 | 741 |
| Omicron | 6168 | 1751 |
| PMS | 1348 | 1554 |
| S2-domain | 4185 | 1274 |
| <u>WT</u> | 7997 | 1731 |

**Table 3.** IL-6 dataset statistics. We only kept the sequences that were unique.

| **Mutant** | **Negatives** | **Positives** |
| --- | --- | --- |
| C72A | 24936 | 461 |
| C78A | 13144 | 530 |
| D162A | 11041 | 953 |
| D168A | 12179 | 280 |
| D54A | 28576 | 494 |
| D99A | 22459 | 437 |
| E108A | 19040 | 626 |
| E138A | 9525 | 1403 |
| E51A | 16834 | 832 |
| E87A | 25580 | 508 |
| F102A | 23287 | 616 |
| F153A | 10255 | 1159 |
| G105A | 12900 | 863 |
| G63A | 27015 | 532 |
| I57A | 28008 | 336 |
| I60A | 27913 | 319 |
| K69A | 26956 | 438 |
| L120A | 12570 | 753 |
| L126A | 13367 | 522 |
| L129A | 12863 | 611 |
| L90A | 11962 | 251 |
| P42A | 17045 | 928 |
| P93A | 20436 | 345 |
| Q144A | 8815 | 1412 |
| Q45A | 16000 | 831 |
| S135A | 15253 | 996 |
| S81A | 24771 | 450 |
| T117A | 14471 | 734 |
| T165A | 12996 | 945 |
| T48A | 17277 | 877 |
| <u>WT</u> | 12433 | 540 |

Each of the sequences was modeled using NanoBodyBuilder2 (Abanades et al. 2023). We assess the fitness of each of the sequences by calculating the perplexity as given by the inverse folding model. Perplexity is a measure of amino acid diversity predicted at each position and reflects the structural constraints of the protein sequence. The model calculates probabilities for each amino acid based on the logits, where probabilities are normalized using a softmax function given by eq. 1

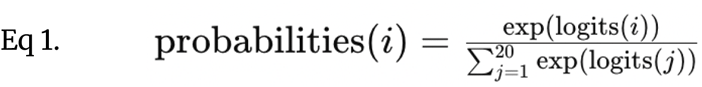

Perplexity is then computed as eq. 2.

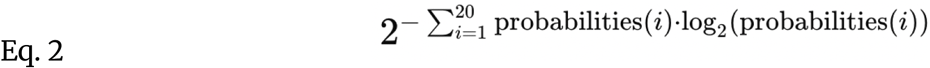

Here, lower values indicate fewer plausible amino acids constrained by the structure. This metric effectively captures how well inverse folding models predict structurally constrained, high-fitness sequences, with potential correlation to high-affinity binders.

### 2.3 Computational oracles for binding prediction

For the sequence design experiments we used computational oracles as proxies for experimental validation. It was shown previously, also on the studies releasing the IL-6 and SARS-COV-2 datasets used here, that using relatively simple neural networks applied to such large datasets one can obtain very good performance. Here we devised a network inspired by those of Mason et al. and Lim et al. (Mason et al. 2021; Lim, Adler, and Johnson 2022).

The study by Mason et al. employed two types of neural networks: long short-term memory recurrent neural networks (LSTM-RNNs) and convolutional neural networks (CNNs). The LSTM-RNNs were chosen for their ability to retain sequential information, making them suitable for correlating sequence order with antigen-binding specificity. In contrast, CNNs were used for their strength in recognizing spatial dependencies in data, utilizing learnable filters to identify key sequence features. Both models were fed input data in the form of one-hot encoded amino acid sequences, where each sequence position corresponded to a specific residue and was represented in a 10 x 20 matrix. The neural networks underwent hyperparameter optimization through grid search methods, and their architectures were validated using k-fold cross-validation. During training, the datasets were split into training and testing sets, with performance evaluated based on metrics such as ROC curves and precision-recall analyses. These models excelled at classifying antigen binders and non-binders with high accuracy and were instrumental in exploring a vast computational sequence space for therapeutic antibodies.

The network of Lim et al. utilized convolutional neural networks (CNNs) for classifying antibody sequences as binders or non-binders to CTLA-4 and PD-1 antigens. Antibody CDR-H3 sequences from both heavy and light chains were encoded into 2D numerical matrices based on BLOSUM substitution scores, with dimensions standardized via gap-padding to accommodate variable sequence lengths. These encoded “images” served as the input for the CNNs. The network architecture included three convolutional layers followed by a dense neural network layer, culminating in a binary classification output. Hyperparameters such as filter numbers, kernel sizes, dropout rates, and dense layer nodes were tuned using randomized search. Performance was assessed with metrics like prediction accuracy, AUC values, and Matthews correlation coefficients, achieving over 91% accuracy. Additionally, generative adversarial networks (GANs) were employed to create synthetic antibody sequences, where a generator network produced sequence-like outputs, and a discriminator network validated their authenticity. This generative approach expanded the repertoire of potential antibody candidates, demonstrating the utility of deep learning in antibody discovery and engineering.

In both cases, the original networks were used for CDR-H3 prediction, but in our datasets, the entire variable region can be used to offer a predictive signal. Therefore we extended the input to cover the entire variable region. Here we only employed the CNN architecture rather than LSTM, for ease of development. We devised a 2-layer CNN network, trained with binary cross entropy loss on the binder non-binder labels. The learning rate was 0.001, using an ADAM optimizer. We strive not to over engineer these networks as we were apprehensive that in deep mutational scanning experiments it is very easy to over-fit.

In both cases, we trained only on the wildtype datasets as we did not want to mix distributions between cognate but distinct mutated antigens. In each case we used nn 8:1:1 training, validation, test split. Training was stopped when validation loss did not show improvement. For the IL-6 dataset, test set accuracy reached 95% whereas for SARS-COV-2 89%, taking a cutoff of arbitrary 0.5 applied to the sigmoid output (where 0 was the negative label and 1 the positive one).

### 2.4. Availability

nanoFOLD weights and usage instructions are available for non-commercial use by non-commercial organisations under the following link.

## 3. Results

### 3.1. Benchmarking nanoFOLD on sequence recovery versus antibody and protein-generic models

Inverse folding methods are trained on an objective to predict a sequence given backbone coordinates. The measure of accuracy here is the ‘recovery rate’, which reflects the degree of agreement of the sequence predicted by a model and the native one associated with the real coordinates.

We performed two recovery tests, one using the crystal structures (Xtal test) with 43 PDBs and model test with 1064 PDBs (Table 4). It was challenging to rule out whether certain crystal structures and models were in the training set of ESM-IF and thus AntiFold. For this reason we reason that our model should do at least as well as ESM-IF on the crystal test set, since we use ESM-IF as basis for fine-tuning. We calculated the recovery on two main regions of interest, the entire variable region (VHH) or just the CDR-H3.

**Table 4.**
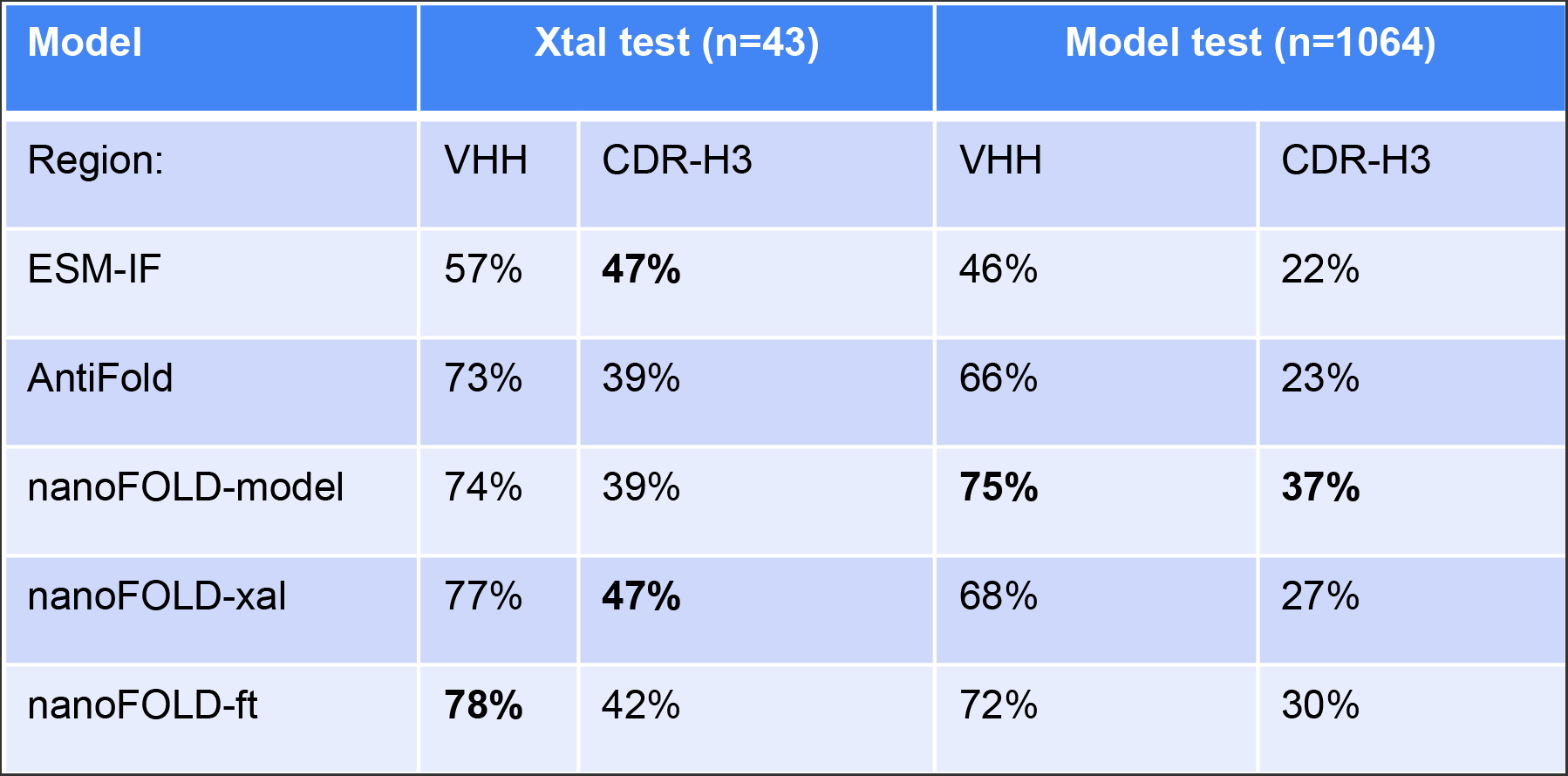
Recovery of nanobody variants using the inverse folding method. We benchmarked ESM-IF, that is the foundation model for inverse folding, AntiFold, antibody-specific fine-tuning of ESM-IF and three variants of nanoFOLD that are nanobody-specific fine tunings of ESM-IF.

| Model | Xtal test (n=43) |  | Model test (n=1064) |  |
| --- | --- | --- | --- | --- |
| Region: | VHH | CDR-H3 | VHH | CDR-H3 |
| ESM-IF | 57% | **47%** | 46% | 22% |
| AntiFold | 73% | 39% | 66% | 23% |
| nanoFOLD-model | 74% | 39% | **75%** | **37%** |
| nanoFOLD-xal | 77% | **47%** | 68% | 27% |
| nanoFOLD-ft | **78%** | 42% | 72% | 30% |

nanoFOLD-based models perform the best across most tests. On the crystal structures dataset, ESM-IF performs worse on the entire variable region, but together with nanoFOLD-xal it gets the highest recovery of CDR-H3. It appears that fine tuning nanoFOLD-model on the crystal dataset, resulting in nanoFOLD-ft did not bring its performance on the CDR-H3 on par with nanoFOLD-xal, but improved on the framework. On the models dataset, nanoFOLD-model outperforms other models by a comfortable margin with 75% for the variable region and 37% for CDR-H3.

It appears that fine-tuning on the crystal dataset (nanoFOLD-ft) did not maintain the best performance on the model dataset, and did not match nanoFOLD-xal on the crystal dataset. Experimentation with more complex training regimens, so as not to run into the issue of catastrophic forgetting (Kirkpatrick et al. 2017) did not yield better results.

Nevertheless, the big picture of fine-tuning ESM-IF on the nanobody dataset, appears to yield positive results, with increased capacity to handle this type of molecule. Nevertheless, recovery by itself is currently of limited interest for engineering and protein design, and thus we tested the model on two experimental datasets for realistic applications.

It was demonstrated that large-scale protein models such as ESM enable prediction of certain protein properties in zero-shot fashion (Meier et al. 2021). We hypothesized that the inverse folding models could behave in a similar fashion, with the higher-fitness space (smaller perplexity, model more confident about sequence-structure-fit) being enriched for molecules of interest, in our case, binders.

## 3.2. Benchmarking nanoFOLD on enriching the NGS dataset for binders in zero-shot fashion

We checked whether employing inverse folding can have a beneficial effect in a commonplace antibody discovery task, NGS-based selections. Here, one is tasked in identifying bindiners from non-binders in a single dataset.

Following work of Hie et al. (Hie et al. 2023), we hypothesized that binders might have propensity for better fitness and thus models such as inverse folding that model such fitness landscapes might offer a way to recover them from the entire dataset (Figure 1A). It must be noted that such methods do not take the molecular complementarity of antibody-antigen into account, rather it would appear that higher fitness sequences correlate with more stable molecules, which translates into enrichment for binders. In case of NGS-based selections, there is no antibody-antigen structure to build upon and thus such antigen-less methods that could enrich binders from a large set of antibodies are desired.

**Figure 1.**
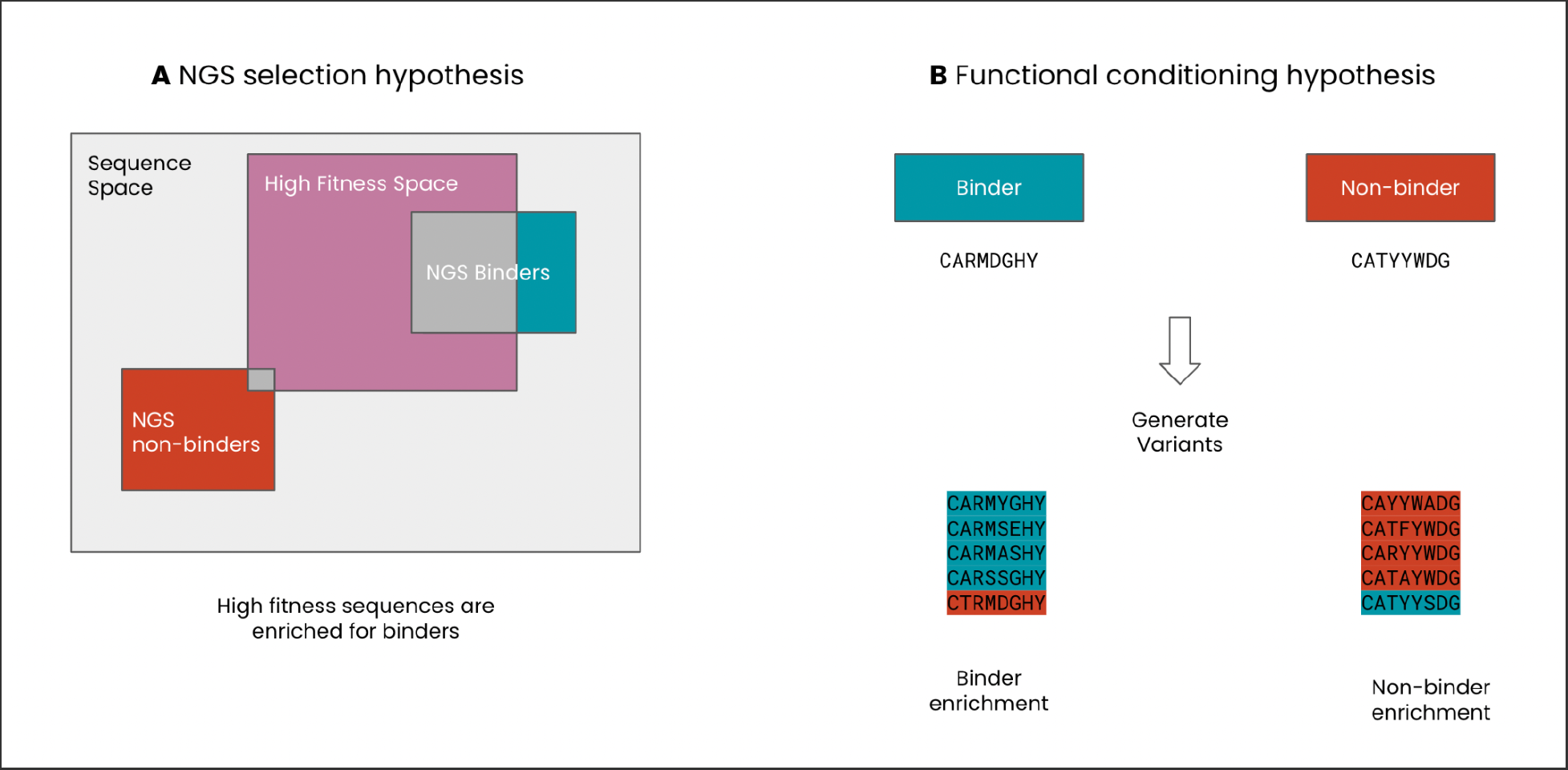
Nanobody design experiments. We tested nanoFOLD models on two nanobody-discovery tasks, NGS-based selections and sequence re-design. **A**. We hypothesized that the inverse folding would define a fitness space, with the more fit sequences being enriched for binding. **B**. We hypothesized that starting with a structure with a predefined function, here binding/nonbinding, the generated sequences with the same fold would largely stay within that prior functional space.

We created a crystal structure of each sequence in the IL-6 and SARS-COV-2 datasets using NanoBuilder2. For each structure, we calculated the perplexity of the sequence that was used to generate the model. The lower the perplexity, the more confident the model is of predictions, theoretically indicating ‘better fitness’.

Within each mutant, binders are a certain proportion that gives a random baseline for enrichment. For instance if there are 5% of binders in any dataset, a method selecting binders at random should average guessing 5% right - surpassing such a percentage is indicative of a model surpassing the random baseline. For each mutant, we looked at the proportion of binders in top 200 picks. This was supposed to simulate a situation where one needs to perform selections from NGS and has roughly 200 clones worth of plate space experimentally. We show the proportion of binders in the top 200 as given by ESM-IF, AntiFold and the three nanoFOLD models for IL-6 (Table 5) and SARS-COV-2 (Table 6).

**Table 5.** IL-6 NGS sorting. For each mutant, we sorted the sequences by the inverse folding scores. The top 200 sequences were picked as ‘final selections’ and the proportion of binders within these were checked. Baseline column gives the proportion of binders in any given dataset - having a value above that is indicative of model enriching the top 200 in binders.

| Variant | esm | antifold | nanofold_xtal_30000 | nanofold_model_95000 | nanofold_ft_137000 | Baseline |
| --- | --- | --- | --- | --- | --- | --- |
| _C72A_ | 0.02 | 0 | 0.04 | 0.03 | **0.07** | 0.02 |
| _C78A_ | 0.02 | 0.01 | 0.03 | 0.03 | **0.1** | 0.03 |
| _D162A_ | 0.02 | 0.01 | 0.03 | 0.06 | **0.11** | 0.05 |
| _D168A_ | 0 | 0.01 | 0.01 | 0.02 | **0.05** | 0.01 |
| _D54A_ | 0.01 | 0.01 | **0.02** | 0.01 | 0.01 | **0.02** |
| _D99A_ | 0 | 0 | 0.01 | 0.01 | **0.03** | 0.02 |
| _E108A_ | 0.01 | 0 | 0.01 | 0.02 | **0.07** | 0.02 |
| _E138A_ | 0.04 | 0.02 | 0.11 | 0.09 | **0.22** | 0.06 |
| _E51A_ | 0.02 | 0.01 | 0.06 | 0.04 | **0.1** | 0.04 |
| _E87A_ | 0 | 0 | 0.01 | 0.01 | **0.02** | **0.02** |
| _F102A_ | 0.01 | 0.01 | 0.02 | 0.01 | **0.05** | 0.02 |
| _F153A_ | 0.03 | 0.01 | 0.09 | 0.09 | **0.15** | 0.05 |
| _G105A_ | 0.02 | 0 | 0.06 | 0.06 | **0.11** | 0.04 |
| _G63A_ | 0.01 | 0 | 0.02 | 0.03 | **0.04** | 0.02 |
| _I57A_ | 0 | 0.01 | 0.01 | **0.02** | 0.01 | 0.01 |
| _I60A_ | 0.01 | 0.01 | **0.02** | 0.01 | **0.02** | 0.01 |
| _K69A_ | 0.01 | 0 | 0.02 | 0.02 | **0.03** | 0.02 |
| _L120A_ | 0.01 | 0.01 | 0.03 | 0.04 | **0.09** | 0.03 |
| _L126A_ | 0.01 | 0.01 | 0.04 | 0.02 | **0.07** | 0.02 |
| _L129A_ | 0.03 | 0.02 | 0.05 | 0.05 | **0.1** | 0.03 |
| _L90A_ | 0 | 0.01 | 0.02 | 0.03 | **0.08** | 0.01 |
| _P42A_ | 0.03 | 0.01 | 0.05 | 0.04 | **0.07** | 0.05 |
| _P93A_ | 0 | 0.01 | 0.01 | 0.01 | **0.02** | 0.01 |
| _Q144A_ | 0.04 | 0.01 | 0.08 | 0.07 | **0.16** | 0.06 |
| _Q45A_ | 0.04 | 0.01 | 0.06 | 0.04 | **0.09** | 0.05 |
| _S135A_ | 0.02 | 0 | 0.04 | 0.03 | **0.08** | 0.03 |
| _S81A_ | 0.01 | 0 | 0.02 | 0.02 | **0.06** | 0.02 |
| _T117A_ | 0.02 | 0.01 | 0.04 | 0.04 | **0.08** | 0.04 |
| _T165A_ | 0.01 | 0.02 | 0.04 | 0.03 | **0.1** | 0.04 |
| _T48A_ | 0.03 | 0.01 | 0.05 | 0.05 | **0.11** | 0.05 |
| _WTs_ | 0.05 | 0.01 | 0.07 | 0.08 | **0.16** | 0.04 |

**Table 6.** SARS-COV-2 NGS sorting. For each mutant, we sorted the sequences by the inverse folding scores. The top 200 sequences were picked as ‘final selections’ and the proportion of binders within these were checked. Baseline column gives the proportion of binders in any given dataset - having a value above that is indicative of model enriching the top 200 in binders.

| Variant | esm | antifold | nanofold_xtal_30000 | nanofold_model_95000 | nanofold_ft_137000 | Baseline |
| --- | --- | --- | --- | --- | --- | --- |
| _Alpha_ | 0.2 | 0.09 | 0.32 | **0.4** | 0.35 | 0.3 |
| _Alpha+E484K_ | 0.31 | 0.07 | 0.44 | **0.51** | 0.47 | 0.36 |
| _Alpha+K417N_ | 0.26 | 0.08 | 0.39 | **0.47** | 0.44 | 0.37 |
| _Beta_ | 0.3 | 0.07 | 0.37 | **0.43** | 0.39 | 0.3 |
| _D614G_ | 0.22 | 0.1 | 0.26 | 0.3 | **0.34** | 0.28 |
| _Delta_ | 0.15 | 0.1 | 0.22 | 0.24 | **0.29** | 0.18 |
| _Kappa_ | 0.5 | 0.38 | 0.56 | **0.62** | 0.6 | 0.48 |
| _Lambda_ | 0.41 | 0.37 | 0.49 | 0.45 | **0.5** | 0.36 |
| _OC43_ | 0.21 | 0.06 | 0.23 | **0.25** | 0.22 | 0.2 |
| _Omicron_ | 0.22 | 0.19 | 0.26 | **0.39** | 0.24 | 0.22 |
| _PMS_ | 0.6 | 0.46 | 0.65 | **0.82** | 0.75 | 0.54 |
| _S2-domain_ | 0.26 | 0.11 | 0.33 | 0.3 | **0.39** | 0.23 |
| _WT_ | 0.11 | 0.03 | 0.18 | **0.28** | 0.24 | 0.18 |

For all mutants, a nanoFOLD-based method surpassed the random baseline. In many cases, AntiFold and ESM-IF failed to surpass the random baseline, which is hard to interpret. The top two performers are nanoFOLD-FT and nanoFOLD-model. The nanoFOLD-FT model surpasses the baseline in 29 out of 31 experiments on IL-6 and 13 out of 13 experiments on SARS-COV-2. The nanoFOLD-model model surpasses the baseline in 13 out of 31 experiments on IL-6 and 13 out of 13 experiments on SARS-COV-2. Performance of nanoFOLD-model on the IL-6 dataset however is questionable as the surpassing of random baseline happens within the limits of statistical significance. On IL-6 dataset, nanoFOLD-model surpassed nanoFOLD-ft only in one instance. On the SARS-COV-2 dataset, nanoFOLD-model surpassed nanoFOLD-ft in 9 out of 13 experiments. The unequal spread, especially in case of SARS-COV-2, indicates that the signal is there as judged by multiple runs on multiple mutants, but the methods are still quite noisy and dataset-specific. The enrichments are not radical, in the order of several percentage points, especially for IL-6. Nevertheless, given the low computational cost of running such an experiment, relative to experimental validation of the binders, the results suggest certain utility that perhaps could be further improved.

The reason for better performance of nanoFOLD models in this context could be the structural modeling function used, NanoBodyBuilder2. AntiFold is based on AbodyBuilder2 whereas pure ESM-IF on AlphaFold2. Therefore, the methods might not receive the coordinates from the generating function they were trained on, which might affect their performance. For equal footing with the original ESM-IF one would need to generate all models using AlphaFold2. Running AlphaFold2 on such a scale would be computationally unreasonable. To have equal footing with AntiFold one would have to generate the single chain models with ABodyBuilder2. It is not clear how to reasonably model single-domain antibodies using Vh/Vl specific ABodyBuilder2.

### 3.3. Generating binders against IL-6 and SARS-COV-2

We hypothesized that inverse folding can generate novel sequences, emulating the function of the parental structure, in this case binding/non-binding (Figure 1B). Performing inverse folding on antibody-antigen complexes was shown to recapitulate binders (Shanker et al. 2023), which is plausible as the novel sequences are given the full molecular interaction context. In a realistic discovery campaign structural complex information is rarely available, especially in the early stages. Sequences with the structural model of a binder are readily available, so generating novel sequence binder based on a known one is desirable. Generating sequences without the knowledge of the interacting partner is challenging, so we checked to what extent the fitness space explored by the inverse folding algorithm of an individual nobody molecule recapitulates the functional space relevant to our discovery case (binding). The remaining issue in our scenario was to check whether the generated nanobodies bind the antigen as we did not have such experimental information.

Several previous studies developed oracles on the basis of binder/non-binder datasets for single targets e.g. HER2 (Mason et al. 2021), CTLA4/PD1 (Lim, Adler, and Johnson 2022), IL6 (Tsuruta et al. 2023), SARS-COV-2 (Tsuruta et al. 2024) and an aggregate of the aforementioned targets (Barton et al. 2024). We repurposed the Mason et al. simple CNN network and trained an oracle on wild types of IL-6 and SARS-COV-2. Employing such computational oracles has been shown to provide results in line with experiments, allowing one a proxy for laborious experimentation (Mason et al. 2021; Lim, Adler, and Johnson 2022).

We ran two in-silico re-design experiments: CDR-H3 redesign and full sequence re-design. We gave any inverse model either a structure of a non-binder or a binder and asked to redesign the entire variable region (VHH mode) or only the CDR-H3 (H3 mode). We generated sequences at temperature 0.1 (not much diversity, should reflect native better) and 1.0 (a lot of diversity, should be more distinct from native).

Since we start with a structure of either a known binder, or non-binder we are biasing the model to provide predictions in the functional area of either binding or not binding the target. Therefore, the sequences for redesign were sampled from the test set of either oracle to avoid overfitting. We called a correct prediction when the sequence generated from either binder/non-binder was predicted in the same class by the oracle.

Results from applying the oracles to the IL-6 and SARS-COV-2 datasets are given in Figure 2 and 3. In each case the PROC AUC is better for the nanoFOLD models than the ESM-IF or antibody-specific AntiFold. All models achieve substantially better performance on predicting just CDR-H3 rather than the entire nanobody molecule. The caveat here can be residual binding from the other CDRs that could be biasing the prediction. Nevertheless, similar bias could be expected in an in vitro experiment, where residual binding from the non-redesigned part still plays a large role in specificity, and the novel CDR-H3 at the very least, maintains the function of the template binder.

**Figure 2.**
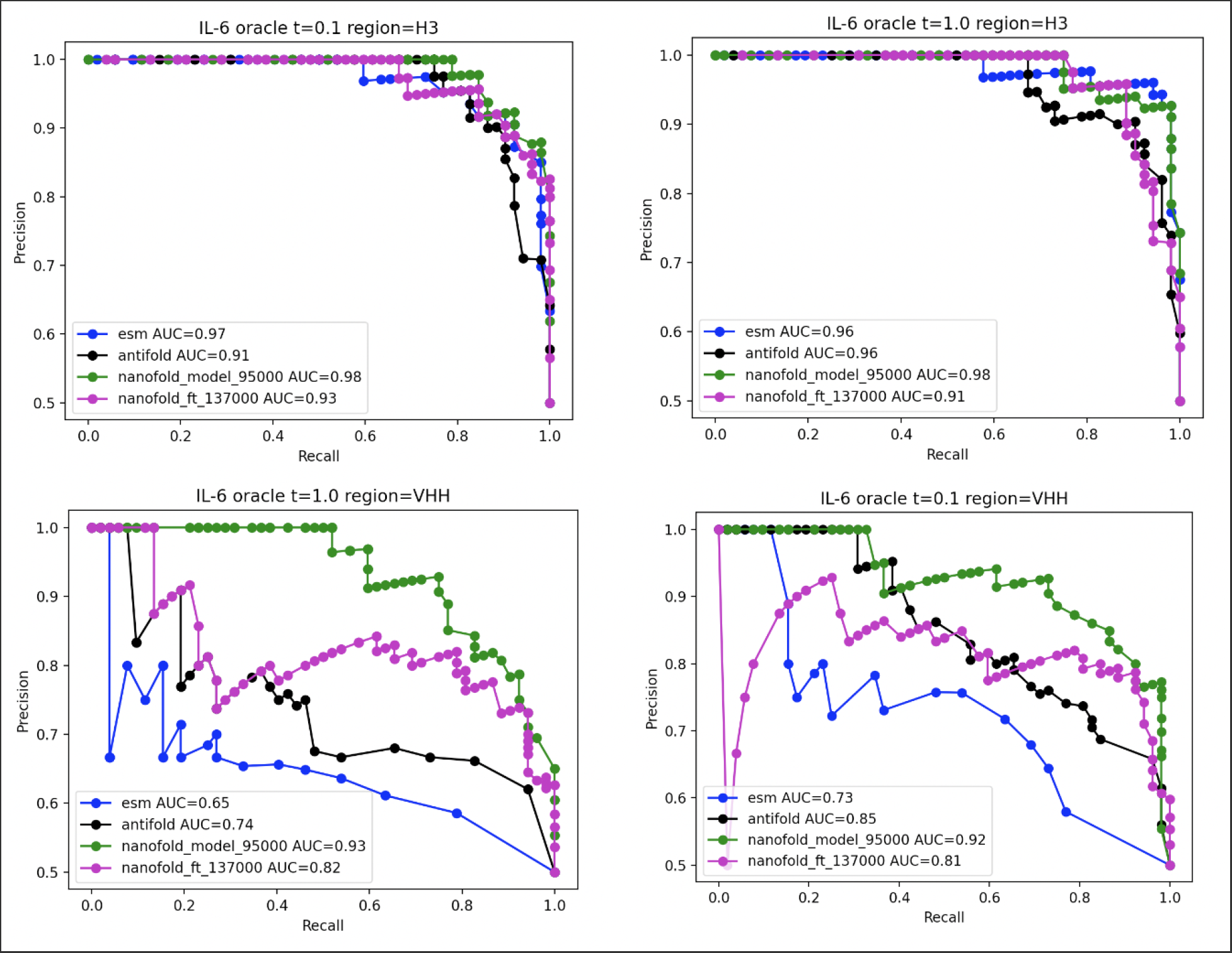
Results of binding oracle experiment on IL-6. We selected binders and non-binders from the wildtype IL-6 cohort. We tasked inverse folding methods to generate either a full variable region sequence (region=VHH) or just the H3 (region=H3). We give the sequence of a binder or non-binder we assume to be conditioning the models to produce binders and non-binders in a few-shot fashion. There was a 50:50 split of binders and non-binders. The sequence variants generated are passed through the binding oracle trained on the IL-6 WT sequences - excluding the set used to generate the sequences.

**Figure 3.**
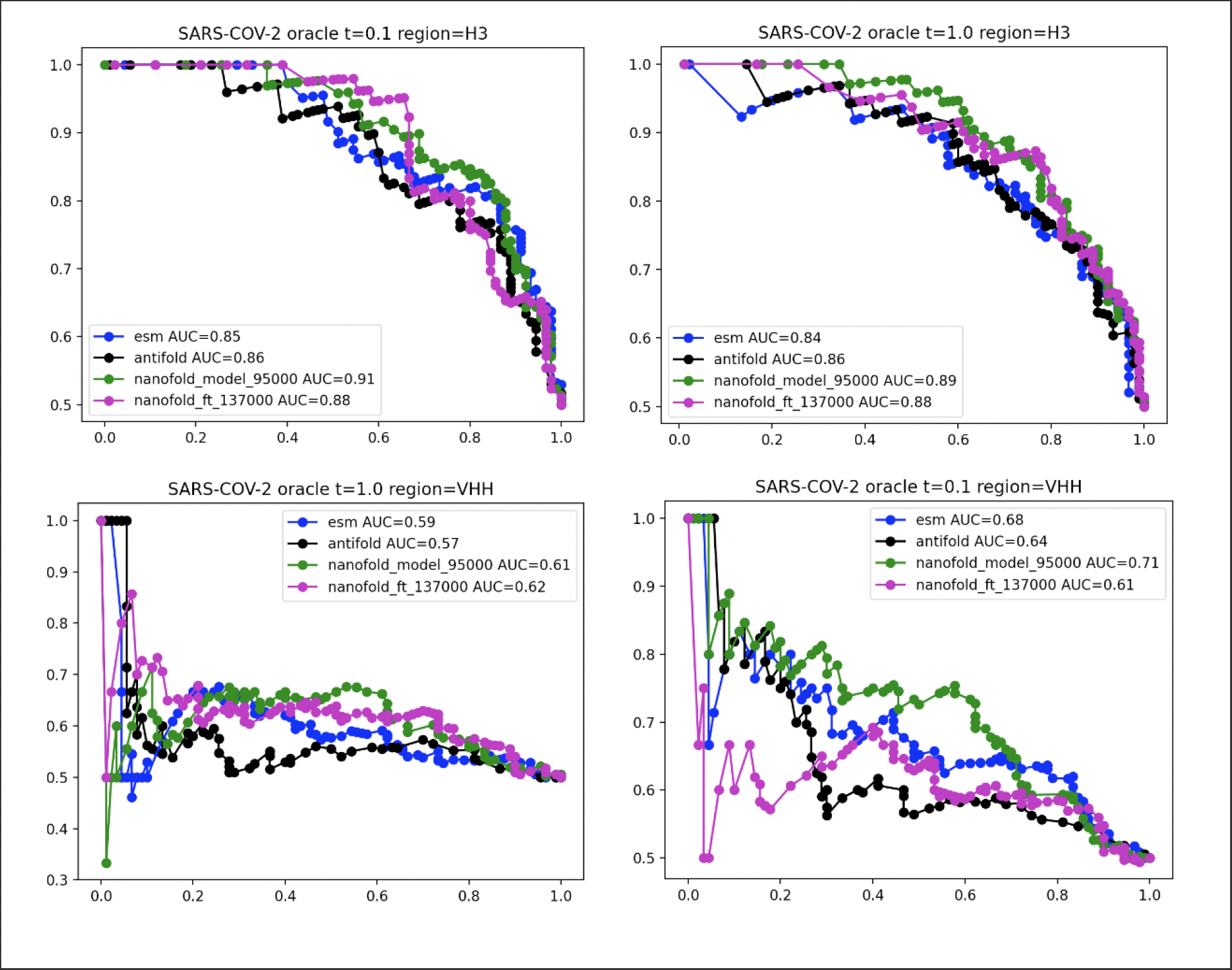
Results of binding oracle experiment on SARS-COV-2. We selected binders and non-binders from the wildtype SARS-COV-2 cohort. We tasked inverse folding methods to generate either a full variable region sequence (region=VHH) or just the H3 (region=H3). We give the sequence of a binder or non-binder we assume to be conditioning the models to produce binders and non-binders in a few-shot fashion.

There was a 50:50 split of binders and non-binders. The sequence variants generated are passed through the binding oracle trained on the SARS-COV-2 WT sequences - excluding the set used to generate the sequences.

## 4. Discussion

Inverse folding has emerged as a viable method for sequence optimization. It offers a solution to the problem of generating viable sequences being able to fit existing coordinates.

Engineering applications range from de novo generation of antibodies (Bennett et al. 2024), through possible affinity optimization (Hummer et al. 2023). Current methods are either protein-general or antibody specific, which dampens the full potential of these methods in applications to specific formats such as nanobodies. Through our development of nanoFOLD weights for ESM-IF we provided a solution for nanobody-specific sequence design using a set of pre-existing coordinates.

We hypothesize that the main limitation in comparisons and applications is the modeling function used for coordinate generation. ESM-IF was trained on AlphaFold2 models whereas AntiFold and nanoFOLD were trained on ABodybuilder2 and NanobodyBuilder2 respectively. Unifying the modeling function is not feasible as AlphaFold2 achieves excellent accuracy on proteins but is doing slightly worse than the antibody-nanobody specific methods. By contrast, the antibody-nanobody specific methods can not achieve the same accuracy on general proteins.

Though the inverse folding methods are chiefly assessed by their ability to perform sequence recovery, it is not the main task that the methods should be benchmarked on. Since we wish to employ these methods for a range of protein optimisation and design applications, the benchmarking cases need to reflect that. Though we performed tests on prior experimental datasets, testing novel binders should ideally be performed in a lab rather than using an in silico oracle alone. Benchmarking sets for antibodies are only coming to view (Chungyoun, Ruffolo, and Gray 2024; Uçar, Malherbe, and Gonzalez 2024). Further development of such benchmark sets for nanobodies is necessary to advance the field of nanobody design.

Altogether, we hope that our novel nanobody-specific inverse folding weights for ESM-IF will help antibody engineers in developing better therapeutics.

## Notes

### Competing Interest Statement

The authors have declared no competing interest.

